# Trapped ion mobility spectrometry (TIMS) and parallel accumulation - serial fragmentation (PASEF) enable in-depth lipidomics from minimal sample amounts

**DOI:** 10.1101/654491

**Authors:** Catherine G. Vasilopoulou, Karolina Sulek, Andreas-David Brunner, Ningombam Sanjib Meitei, Ulrike Schweiger-Hufnagel, Sven Meyer, Aiko Barsch, Matthias Mann, Florian Meier

**Affiliations:** Max-Planck Institute of Biochemistry, Martinsried, Germany; NNF Center for Protein Research, Copenhagen, Denmark; PREMIER Biosoft, Indore, India; Bruker Daltonik GmbH, Bremen, Germany

## Abstract

Lipids form a highly diverse group of biomolecules fulfilling central biological functions, ranging from structural components to intercellular signaling. Yet, a comprehensive characterization of the lipidome from limited starting material, for example in tissue biopsies, remains very challenging. Here, we develop a high-sensitivity lipidomics workflow based on nanoflow liquid chromatography and trapped ion mobility spectrometry. Taking advantage of the PASEF principle (Meier *et al.*, PMID: 26538118), we fragmented on average nine precursors in each 100 ms TIMS scans, while maintaining the full mobility resolution of co-eluting isomers. The very high acquisition speed of about 100 Hz allowed us to obtain MS/MS spectra of the vast majority of detected isotope patterns for automated lipid identification. Analyzing 1 uL of human plasma, PASEF almost doubled the number of identified lipids over standard TIMS-MS/MS and allowed us to reduce the analysis time by a factor of three without loss of coverage. Our single-extraction workflow surpasses the plasma lipid coverage of extensive multi-step protocols in common lipid classes and achieves attomole sensitivity. Building on the high precision and accuracy of TIMS collisional cross section measurements (median CV 0.2%), we compiled 1,327 lipid CCS values from human plasma, mouse liver and human cancer cells. Our study establishes PASEF in lipid analysis and paves the way for sensitive, ion mobility-enhanced lipidomics in four dimensions.

---

Every cell contains large amounts of lipids in various concentrations and chemical compositions^1,2^. Aberrant lipid homeostasis is a hallmark of many diseases, including cancer^3,4^ and metabolic disorders^5^. Therefore, analyzing lipidomes on a large scale promises novel insight into basic biology, as well as the onset and progression of disease^6–8^.

Lipid extracts from biological sources can be analyzed either directly via high-resolution mass spectrometry^9,10^ or via online liquid chromatography (LC-MS)^8^. Lipids can be identified based on accurate mass and the MS^2^ or MS^3^ fragmentation pattern, which is facilitated by recent software developments and ever growing reference databases^11–14^. Established LC-MS lipidomics workflows separate lipids at flow rates in the higher micro- or milliliter per minute range, which ensures high sample throughput and robustness, but also compromises sensitivity. As the available sample amount becomes a limiting factor, for example with small tissue sections from biobanks or small cell sub-populations, it is increasingly attractive to employ nanoflow chromatography^15–17^.

MS technology has greatly improved and state-of-the-art high-resolution Orbitrap or time-of-flight (TOF) instruments transmit ions very efficiently and achieve low- to sub-ppm mass accuracy^18,19^. The high acquisition speed of TOF analyzers makes them compatible with very fast separation techniques such as ion mobility spectrometry (IMS)^20,21^. Nested in-between LC and MS, IMS provides an additional dimension of separation based on the ions’ shape and size (collisional cross section, CCS). This is particular interesting for lipidomics, as it provides an opportunity to separate otherwise unresolved isomers^22–25^. Furthermore, the chemical structure of lipids is closely linked to the CCS, which allows predictions by machine learning and could facilitate lipid identification^26–29^.

Trapped ion mobility spectrometry (TIMS) is a relatively new form of IMS that inverts the separation principle of classical drift tube ion mobility^30–34^. Ions entering the TIMS analyzer are positioned in an electrical field by the drag of a gas flow. Lowering the electrical force releases ions from the TIMS device separated by their ion mobility, while the IMS resolution is proportional to the user-defined ramp time. It can be tuned to over 200 CCS/ΔCCS, for example to separate isomeric lipids with distinct double bond positions or geometries^35^. We recently introduced a MS scan mode termed parallel accumulation serial fragmentation (PASEF) that synchronizes TIMS with MS/MS precursor selection^36^. In proteomics, PASEF increases MS/MS scan rates more than ten-fold, importantly, without the loss of sensitivity that is otherwise inherent to faster acquisition rates^37^.

Here, we explore whether the PASEF principle can be transferred to lipidomics. We build on nanoflow chromatography to establish a rapid PASEF lipidomics workflow capable of comprehensively analyzing low sample amounts. To investigate the potential of the additional TIMS dimension, we set out to compile a high-precision lipid CCS library from body fluids, tissue samples and human cell lines.

## RESULTS

### Development of the nanoflow PASEF lipidomics workflow

We aimed to develop a rapid workflow that enables global lipid analysis in a straightforward manner (**Fig. 1**). Our lipid extraction protocol is applicable to common biological sample types, such as body fluids, tissue, as well as cell lines (**Fig. 1a**) and requires only a few manual liquid handling steps that could easily be automated in the future. We found that our extraction protocol scales well from small sample volumes (1 uL blood plasma) to relatively large cell counts (0.5 million HeLa cells) and can be performed in less than 1 hour.

**Figure 1.**
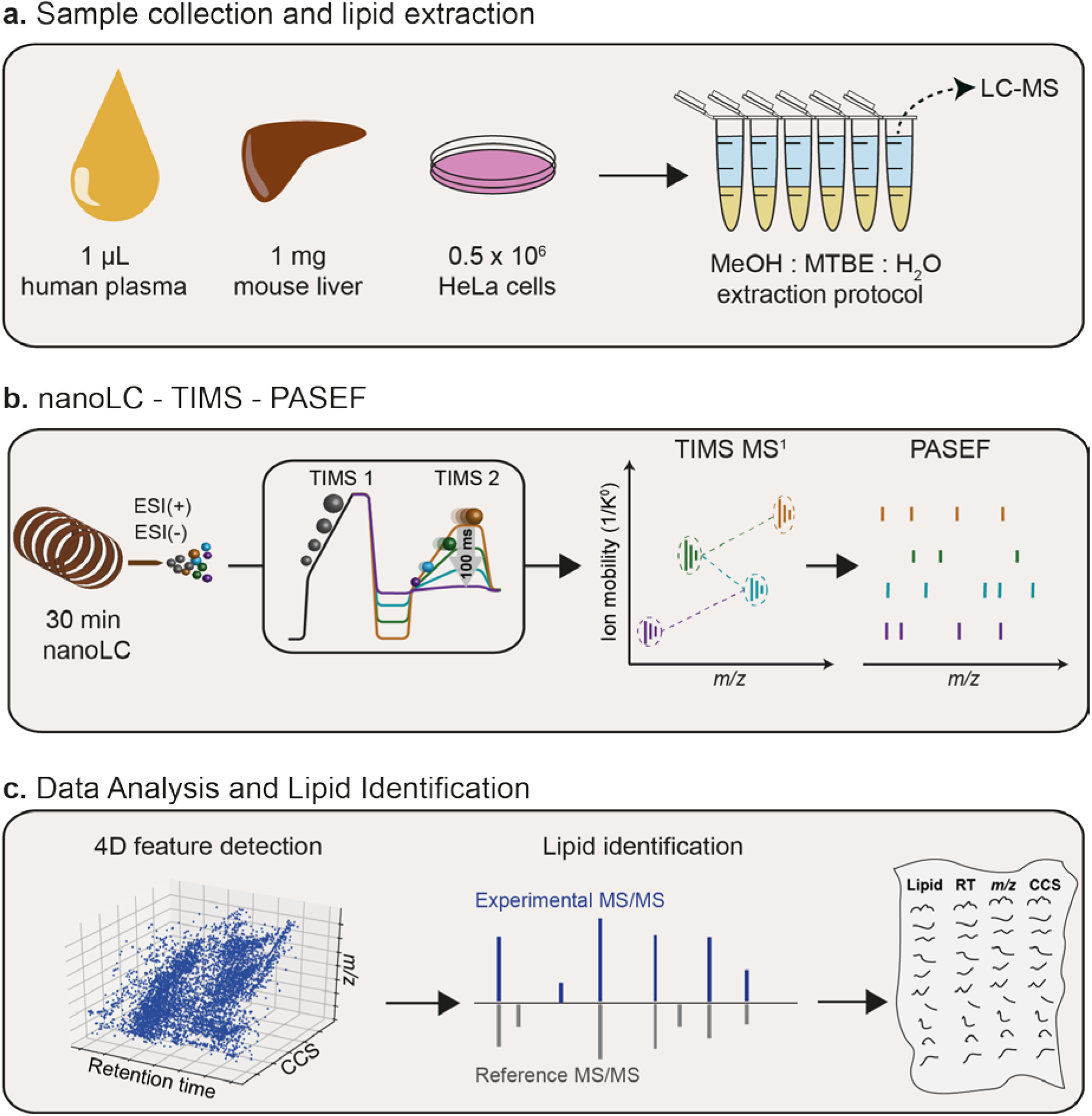
Nanoflow lipidomics with trapped ion mobility spectrometry. **a**, Lipids from various biological sources, such as body fluids, tissues and cells, are analyzed using a single MeOH:MTBE extraction. **b**, The crude extract is injected into a nanoflow liquid chromatography (LC) system coupled online to a high-resolution TIMS quadrupole time-of-flight mass spectrometer (timsTOF Pro). In the dualTIMS analyzer, ions are accumulated in the front part (TIMS 1) while another batch is released as a function of ion mobility from the TIMS 2 analyzer. PASEF synchronizes precursor selection and ion mobility separation, which allows fragmenting multiple precursors in a single TIMS scan at full sensitivity. **c**, Features are extracted from the four-dimensional (retention time, *m/z*, ion mobility, intensity) space and assigned to PASEF MS/MS spectra for automated lipid identification and compilation of comprehensive lipid CCS libraries. *MeOH* = methanol, *MTBE* = methyl-tert.-butyl ether, *CCS* = collisional cross section.

We loaded the lipid extracts directly onto a C_18_ column and eluted them within 30 min, for a total of about one hour total analysis time per sample when using both positive and negative ionization modes (**Fig. 1b**). Chromatographic peak widths were in the range of 3 to 6 s full width at half maximum (FWHM), at least two orders of magnitude slower than TIMS ion mobility analysis (100 ms) and the acquisition speed of high-resolution TOF mass spectra (∼100 µs). As the timsTOF Pro mass spectrometer (Bruker Daltonics) features a dualTIMS analyzer, it allows to use up to 100% of the incoming ion beam^32^ **(Methods**). In this mode, ions are accumulated in the first TIMS analyzer while another batch of ions is separated by ion mobility in the second TIMS analyzer. TIMS closely resembles classical drift tube ion mobility, however, ions arrive at the mass analyzer in the inverse order, i.e. low-mobility (and high-mass) ions are released first, followed by ions with higher mobility (and lower mass). In our experiments with 100 ms TIMS scan time, the ion current accumulated during 100 ms was concentrated into ion mobility peaks of 2-3 ms FWHM, which should lead to a 50-fold increase in signal-to-noise as compared to continuous acquisition. The ion mobility-resolved mass spectra result in two-dimensional heat maps, from which suitable precursor ions are selected for fragmentation in data-dependent MS/MS mode (**Fig. 1b**). With PASEF, multiple precursors are fragmented in each TIMS ramp by rapidly switching the quadrupole (see below). As the collision cell is positioned after the TIMS analyzer in the ion path, fragment ions occur at the same ion mobility position as their precursor ions in MS1 mode.

We make use of this information in the post-processing (**Fig. 1c**) to connect the MS/MS spectra to the corresponding MS features extracted from the four-dimensional data space (retention time, m/z, ion mobility, intensity). Finally, we rely on established lipidomics software to automatically assign lipids to the spectra based on diagnostic fragment ions and database matching (**Methods**). Our workflow automatically converts ion mobility to CCS values and records them for all detected MS1 features and thus all identified lipids.

### Evaluating PASEF in lipidomics

The central element of our workflow is the PASEF acquisition method. PASEF takes advantage of the temporal separation of ions eluting from the TIMS device to select multiple precursors for MS/MS acquisition in a single TIMS scan ^36^. To illustrate, **Figures 2a and b** show representative MS1 heat maps of co-eluting lipids from an LC-TIMS-MS analysis of human plasma. The ions are widely spread in *m/z* vs. ion mobility space, while higher mass roughly correlates with lower ion mobility. In standard MS/MS mode, the quadrupole isolates a single precursor mass throughout the entire TIMS scan time (red diamond in **Fig. 2a**). However, the targeted precursor completely elutes during about 6 out of 100 ms, and thus over 90% of the acquisition time is effectively not used. In PASEF mode, the quadrupole instead switches its mass position within approximately 1 ms to capture as many precursors as possible (red diamonds in **Fig. 2b**). In this example, 10 precursors were selected during a single PASEF scan, which translates into a 10-fold increased MS/MS acquisition rate of about 100 Hz. Importantly, this does not come at a loss in sensitivity because the full precursor ion signal from 100 ms accumulation time is captured.

**Figure 2.**
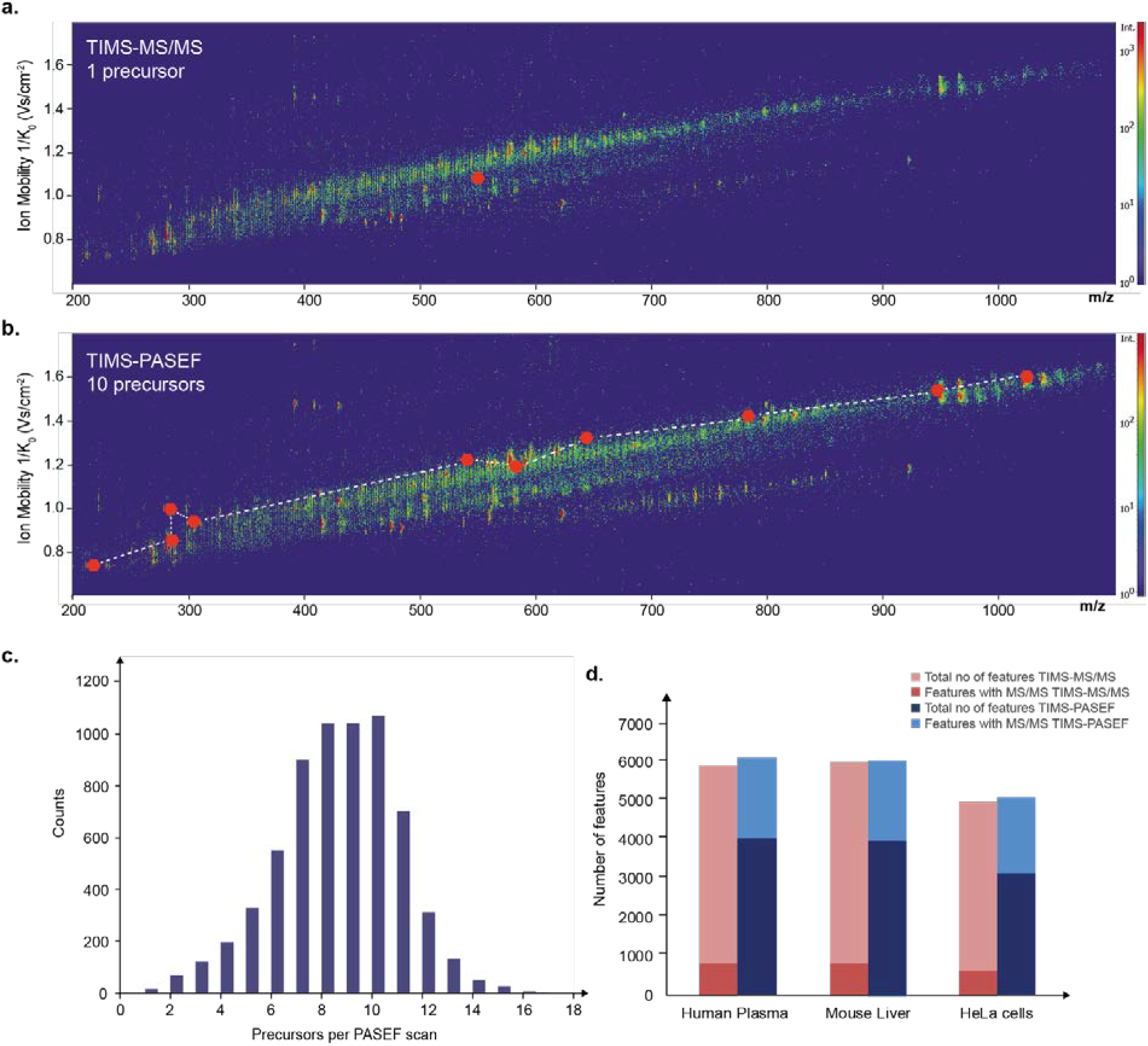
Evaluating PASEF in lipidomics. **a**,**b**, Heat-map visualization of a representative trapped ion mobility resolved mass spectrum of human plasma. Red dots indicate precursors selected for MS/MS fragmentation in the subsequent 100 ms PASEF scan in a, standard TIMS-MS/MS mode and b, TIMS-PASEF mode. **c**, Distribution of the number of precursors per PASEF scan in an LC-MS analysis of human plasma lipid extract. **d**, Total number of 4D features extracted from 30 min runs of human plasma (n=4), mouse liver (n=5) and human cancer cells (n=5) in positive ion mode without (TIMS-MS/MS, red) and with PASEF (TIMS-PASEF, blue). The fraction of features with assigned MS/MS spectra is indicated by a darker color.

In a 30 min analysis of plasma, we found that on average nine precursors were fragmented per PASEF scan (**Fig. 2c**), confirming that the PASEF principle is transferable to lipidomics. In total, we acquired 111,122 MS/MS spectra in 30 min, 8.5-fold more than without PASEF. This fragmentation capacity greatly exceeds the number of expected lipids and, in principle, allows to acquire MS/MS spectra for every suitable isotope pattern detected in a single lipidomics LC run. Here we chose to acquire each PASEF scan twice using two different collision energies to increase the fragment ion coverage of each precursor (**Methods**).

We evaluated the performance of our PASEF method with lipid extracts from human plasma, mouse liver and HeLa cells (**Fig. 2d and Suppl. Fig. 1**). In all three samples types, the 4-dimensional feature detection yielded 4,200 to 6,300 MS1 features above the intensity threshold and after collapsing multiple adducts. In standard TIMS-MS/MS mode, on average 14% of these were fragmented. This fraction increased up to 6-fold with PASEF and, in both negative and positive ionization mode, about two-thirds of all features had corresponding MS/MS spectra. Overall, PASEF almost doubled the number of identified lipids across all runs (**Suppl. Fig. 2**). To further investigate whether PASEF is fast enough to acquire MS/MS spectra of close to all informative lipid features in a short time, we extended the LC gradients to 60 and 90 min (**Suppl. Fig. 3**). Indeed, 90% of all lipids identified with the three times longer gradient were already identified in the 30 min PASEF run, which confirms our hypothesis and suggests that even shorter runtimes could be explored.

### Comprehensive and accurate lipid quantification

Having ascertained that PASEF achieves a very high MS/MS coverage of lipidomics samples, we investigated our automated data analysis pipeline in more detail (**Fig. 3a**). Starting from the thousands of 4D features detected in all replicate injections of human plasma, mouse liver or HeLa cells, we kept those with assigned MS/MS scans for further analysis. PASEF spectra are resolved by ion mobility and the software extracts the MS/MS spectra at the ion mobility position of the respective precursor ion. We then searched all MS/MS spectra considering four lipid categories with the respective lipid classes and subclasses (**Methods**). This yielded 346 to 571 annotations for each sample and ionization mode. We manually inspected all annotations to remove potential false positives based on the reported score and the observed fragmentation pattern. Finally, we grouped co-eluting peaks that were separated by their ion mobility but could not be distinguished based on their MS/MS spectra. However, we kept separate lipids with same MS/MS-based annotation but eluting in close proximity as these are potential isomers. Removing duplicates resulted in 229 to 385 identified unique lipids per experiment. Combining both ionization modes, we identified 437 unique lipids from the equivalent of 0.05 μL plasma per injection, 553 unique lipids from 10 μg liver tissue and 575 unique lipids from approximately 2,000 HeLa cells, with a median absolute precursor mass accuracy of 1.7 ppm **(Suppl. Table S1-S3).** The identified lipid species covered all major lipid classes such as glycerophospholipids (PC, PE, PA, PS, PI, PG), oxidized glycerophospholipids, monoacyl-, diacyl- and triacyl-glycerols, sterol lipids, ceramides, glycosphingolipids and phosphosphingolipids.

**Figure 3.**
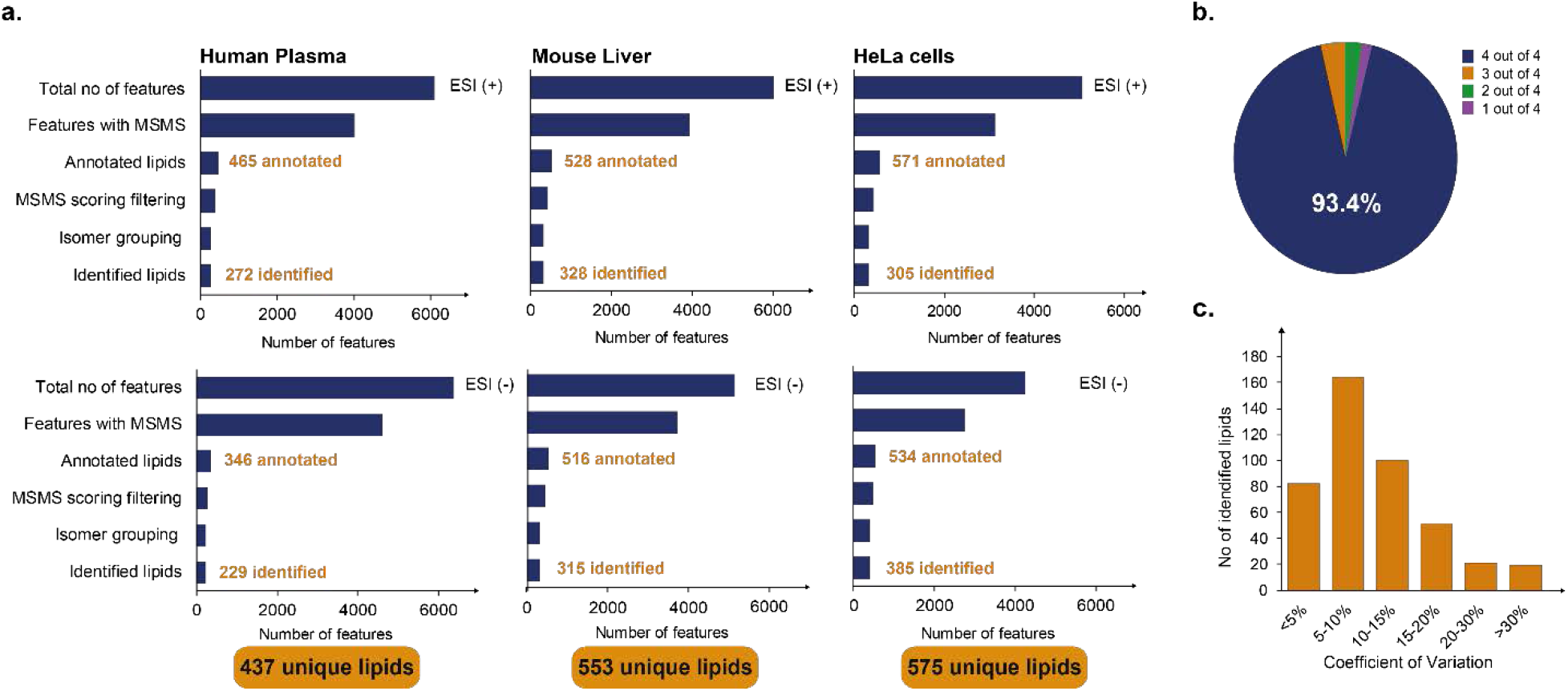
Lipid identification and quantification. **a**, Sequential data analysis steps from the total number of detected 4D features to unique lipids for human plasma, mouse liver and human cancer cells in both ionization modes. **b**, Fraction of lipids quantified in *N* out of five replicate injections of plasma. **c**, Coefficients of variation for 437 lipids label-free quantified in at least two out of four replicate injections of plasma.

In biological or clinical studies, quantitative accuracy is at least as important as cataloguing the lipid composition and may be compromised if lipids are sparsely detected across samples. We hypothesized that the speed of PASEF and the improved signal-to-noise of TIMS should lead to very reproducible label-free quantification. Indeed, we observed that 408 out of the total 437 identified lipids in human plasma were quantified in four out of four replicates (**Fig. 3b**), resulting in a data completeness of 97%. The median coefficient of variation (CV) was 9.2%, and 91% of all quantified lipids had a CV below 20% (**Fig. 3c and Suppl. Table S4**).

From these results, we conclude that our nanoflow PASEF lipidomics workflow covers the lipidome comprehensively with very high quantitative accuracy, while requiring only minimal sample amounts.

### Coverage of the human plasma lipidome

To assess the lipid coverage of our PASEF workflow in more detail, we analyzed the human plasma Standard Reference Material (SRM 1950), a pool from 100 individuals from the United States in the age range of 40 to 50 years, provided by the National Institute of Diabetes and Digestive and Kidney Diseases (NIDDK) and the National Institute of Standards (NIST)^38^. This sample has served as a reference for many lipidomics studies, establishing a range of detectable lipid species and their absolute concentrations^39^. The LIPID MAPS consortium recently compiled consensus results from 31 laboratories, each of which followed their in-house analysis workflow (Bowden *et al.*^40^). In an effort to disentangle the human plasma lipidome, Quehenberger *et al.*^41^ employed class-specific analysis strategies to quantify about 500 lipids from more than 1 mL NIST SRM 1950 plasma.

Taking these two studies as a reference, we first compared the number of identified lipids in each lipid category based on the short name annotation (**Fig. 4a and Suppl. Table S5-S6**). Starting from 1 μL plasma and with a single extraction protocol, we identified on average 76.2% and 71.2% of all glycerolipids and glycerophospholipids reported in both studies, respectively. Remarkably, the total number of lipids in these two abundant plasma lipid categories were 85.1% and 126.0% higher than in the Bowden study, and 89.1% and 81.6% higher than in the Quehenberger study. We observed slightly fewer sphingomyelins, however, still with a very high overlap of 75.8% with both reference studies on average. Analysis of ceramides typically requires specific extraction methods and this category was therefore less comprehensively covered in ours (11.5%) as well as in the Bowden study (22.8%) relative to the class-specific analysis by Quehenberger *et al.* We observed a similar trend for sterol lipids, however, to a lesser extent (54.5%).

**Figure 4.**
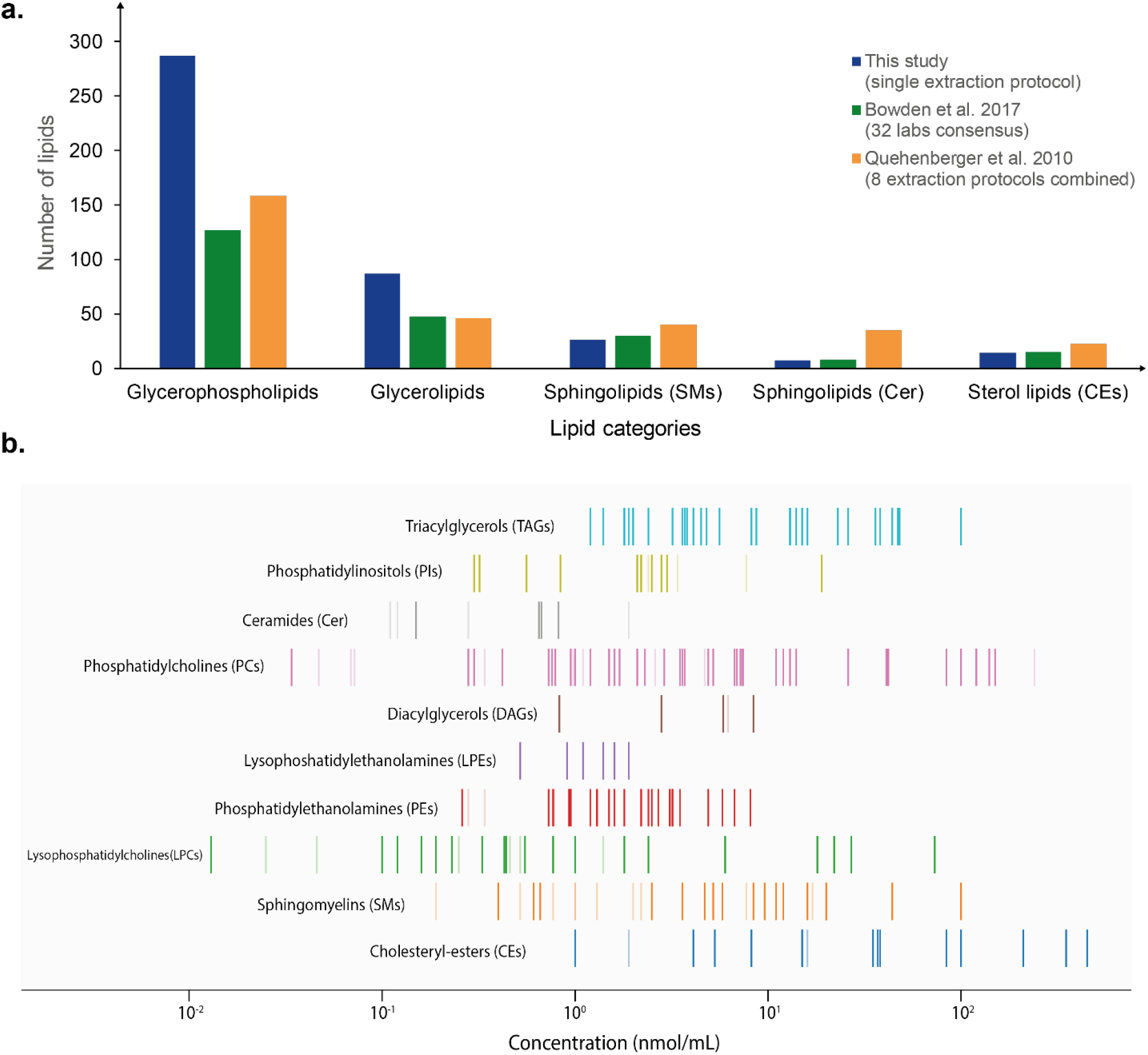
Analysis of 1 µL NIST SRM 1950 human plasma. **a**, Number of identified lipids from major lipid classes in this study and two reference studies from the same standard material^40,41^. **b**, Mapping of lipids identified with our PASEF lipidomics workflow to absolute plasma concentrations reported in ref. 40. Vertical lines indicate the abundance range of reported lipids from different lipid classes and dark color indicates commonly identified lipids in both studies.

To further investigate the sensitivity of our method, we mapped all identified lipids onto absolute plasma concentrations reported in ref. 41 (**Fig. 4b and Suppl. Table S6**). We quantified about 80% of the lipids covering the full abundance range from about 0.01 nmol/mL up to 1,000 nmol/mL. For example, we achieved full coverage of the triacylglycerols and also quantified less abundant lipids such as phosphatidylethanolamines comprehensively. Even though coverage was sparser in the lowest abundance range, we quantified the least abundant lysophosphatidylcholine (LPC 22:1) with a reference concentration of 0.013 nmol/mL. Since we injected only 1/20^th^ of the lipids extracted from 1 μL plasma in each replicate, this translates into a sensitivity in the attomole range for the entire workflow.

### Accuracy and precision of online lipid ^TIMS^CCS measurements

In addition to generating MS/MS spectra for almost all detectable precursors with PASEF, TIMS measures the ion mobility of all identified and unidentified lipids. Because they have the same underlying physics, TIMS ion mobilities can be directly related to drift tube experiments^33,34,42^. We calibrated all TIMS measurements to reduced ion mobility values using well-characterized phosphazine derivatives and converted them to collisional cross sections (^TIMS^CCS) using the Mason-Schamp equation^32,43^.

First, we investigated all 6,100 features detected in the NIST SRM 1950 plasma in positive mode, regardless of their identification as a lipid. Plotting their ^TIMS^CCS values across repeated injections revealed an excellent reproducibility with a Pearson correlation coefficient of 0.998 and a median absolute deviation of 0.17% (**Fig. 5a**). This also holds true in the tissue and cell line experiments, as well as in negative mode (**Suppl. Fig. 4**). Considering only identified lipids, we found a median CV of ion mobilities of 0.30% in repeated injections of the same sample (**Fig. 5b**) and, reassuringly, we achieved this very high precision also across commonly identified lipids from the three different biological samples (**Fig. 5c and Suppl. Table 1-S7**).

**Figure 5.**
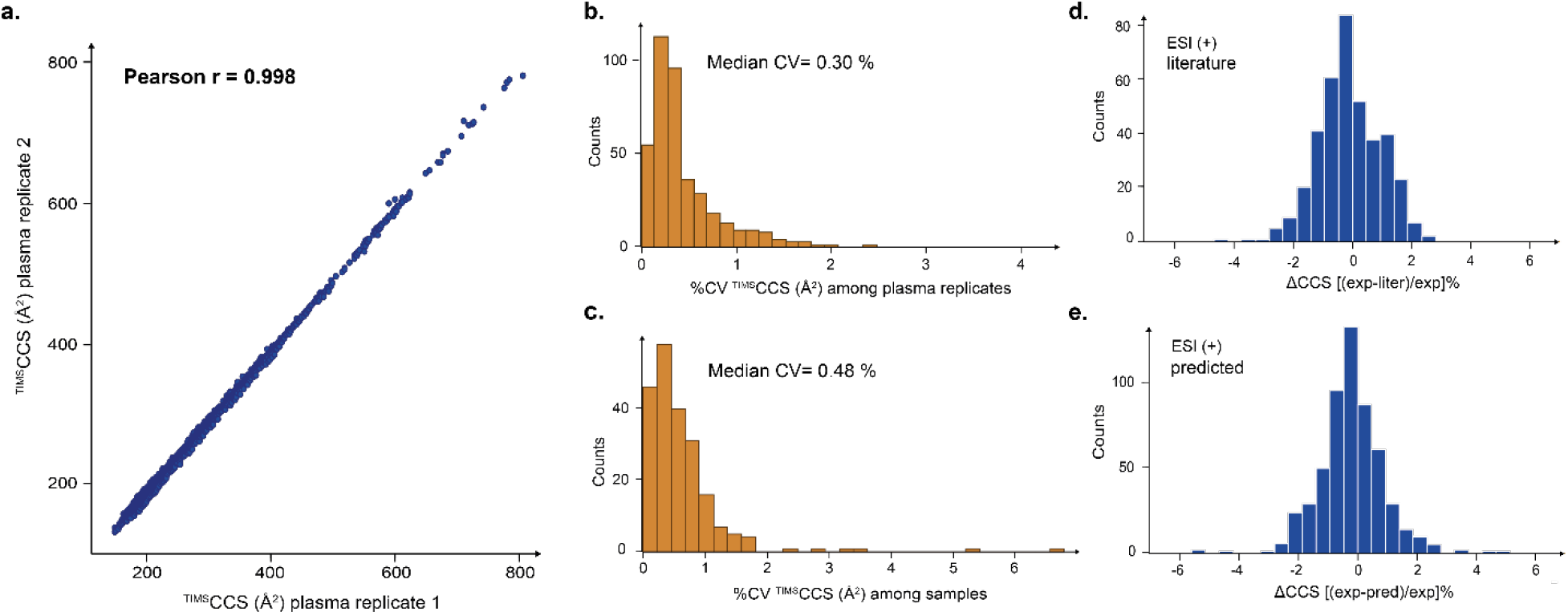
Precise and accurate determination of lipid ^TIMS^CCS values. **a**, Pearson correlation of ^TIMS^CCS values of 6,100 4D features detected in two replicate injections of a human plasma lipid extract. **b**, Coefficients of variation (CV) of ^TIMS^CCS values for lipids commonly identified in replicate injections of the same sample (plasma) and **c**, three different samples (plasma, liver, HeLa). **d**, Relative deviation of experimental ^TIMS^CCS values in this study from literature reports^26^ and **e**, machine learning predictions^27^.

Having ascertained highly reproducible ^TIMS^CCS measurements, we next investigated the accuracy of our results by comparing it to different methods and instrumentation. Our dataset shared 288 and 97 lipid identifications (based on the short name annotation) with recent reports from the Zhu^27^ and McLean^26^ laboratories, which both employed drift tube ion mobility analyzers to establish high-precision reference data. This comparison revealed a very high correlation (r = 0.993) and 95% of all values were within ±2% deviation centered at zero (**Fig. 5d and Suppl. Table S8**). The median absolute deviation was 0.7%, which is in the range of recently reported inter-laboratory variability for standard compounds measured with a commercial drift tube analyzer^44^.

The reproducibility of lipid ion mobility measurements makes them also very attractive for machine learning approaches. The Zhu laboratory has developed a support vector regression model that predicts lipid CCS values from SMILES structures^27^ and which is implemented in the Bruker MetaboScape software (**Methods**). Even though the model was trained on independent data from a different instrument type, predicted and experimental ^TIMS^CCS values correlated very well (r = 0.993) and the relative deviations were normal distributed around zero for lipid classes that were contained in the training data (**Fig. 5e and Suppl. Table S9**). The average absolute deviation was 0.57% and based on the experimental precision demonstrated above, we expect that machine learning models trained direly on TIMS data yield even more accurate predictions.

### The TIMS lipidomics landscape

Data generated by our TIMS lipidomics workflow span a three-dimensional data space in which each feature is defined by retention time, *m/z* and CCS, with intensity as a 4^th^ data dimension. To explore this data space, we compiled all measurements from human plasma, mouse liver and HeLa cells acquired in both ionization modes. The total dataset comprises 1,327 CCS values of 926 unique lipids, representing the four major lipid categories and 15 lipid classes (**Suppl. Table S10**). To make the dataset fully accessible, we provide Suppl. Tab. 1 in a format that follows the standard lipid nomenclature guidelines by the LIPID MAPS consortium^45^ and Lipidomics Standards Initiative (LSI) (https://lipidomics-standards-initiative.org/).

**Figure 6a** shows a three-dimensional representation of all identified lipids color-coded by their respective classes. Each lipid class occupies a discrete space in the conformational landscape, which reflects the structural differences in their chemical composition. Hydrophilic lipids, such as monoacyl- and low molecular weight diacylglycerophospholipids (*m/z* 400-600) that elute first in reversed-phase chromatography, distribute in the CCS dimension from 204 to 253 Å^2^. The second half of the LC gradient is dominated by the large population of glycerolipids and glycerophospholipids, which are often co-eluting, but distinct in mass and CCS. For example, diacylglycerols (DAG) and triacylglycerols (TAG) differ by one acyl chain and also occupy a different CCS space shifted by 54.7 Å^2^. Similarly, the head groups can strongly influence the ion mobility of lipids, as exemplified by PIs and PGs with the same acyl chain composition **(Fig. 6a and Suppl. Fig. 4).**

**Figure 6.**
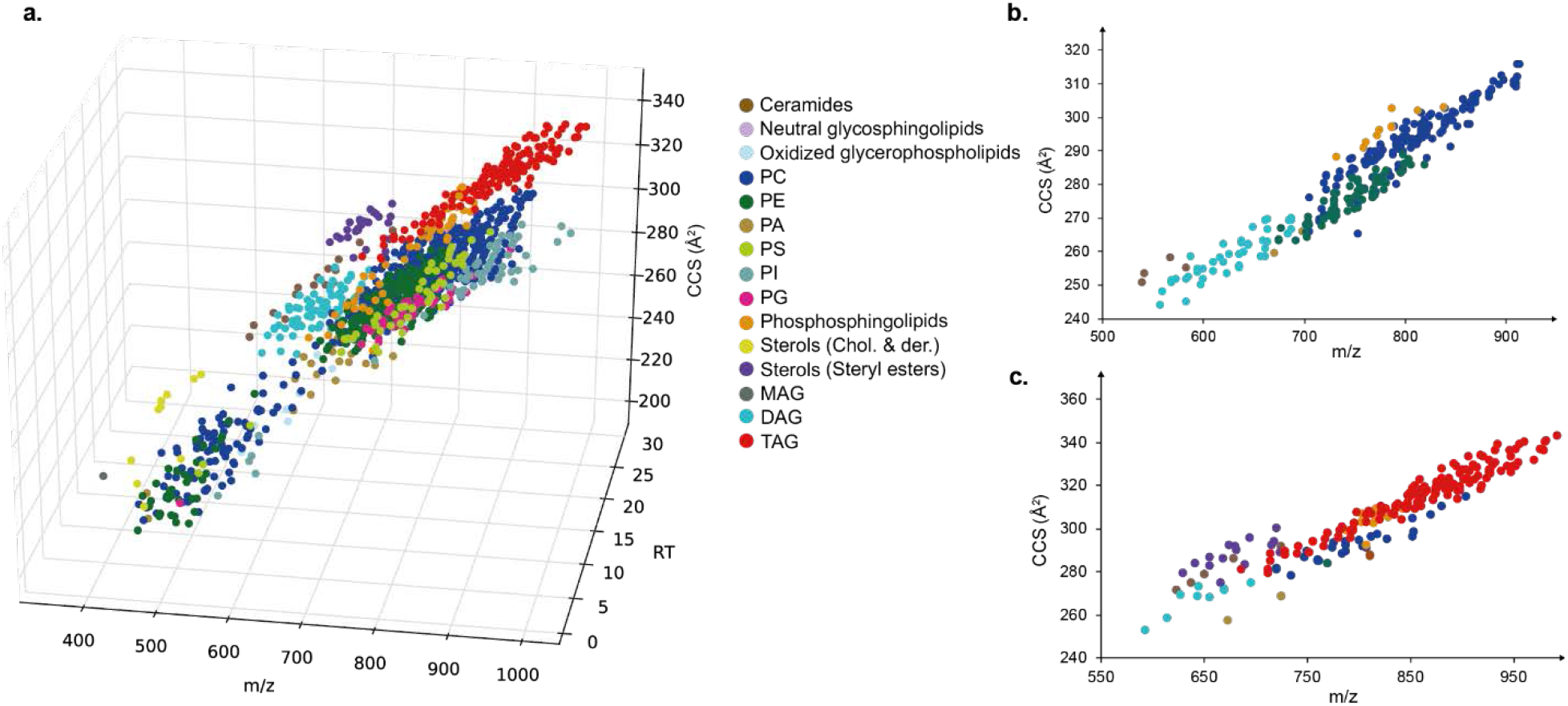
The conformational landscape of lipid ions. **a**, Three-dimensional (RT, *m/z*, CCS) distribution of 1,327 lipids from various classes from three biological samples (plasma, liver, HeLa). **b**, CCS vs *m/z* distribution of lipids eluting between 20 and 23 min. **c**, CCS vs *m/z* distribution of lipids eluting between 24 and 27 min. *PC= Phospatidylcholine, PE=Phospatidylethanolamine, PA= Phosphatidic acid, PI= Phospatidylinositol, PG= Phospatidylglycerol, PS=Phospatidylserine, MAG=Monoacylglycerol, DAG=Diacylglycerol, TAG=Triacylglycerol.*

The investigation of the correlation of lipids mass and ion mobility has been a long term interest in ion mobility spectrometry-based lipidomics^26,27^. TIMS and PASEF provide a very efficient way to extend the scope of such studies to complex biological samples. To illustrate, we zoomed into narrow elution windows of the LC gradient (**Figs. 6b and c)**. In addition to coarse separation of lipid classes, there is a fine-structure within each of them. For example, extension of the acyl chain of phosphocholines (PCs) increases the CCS incrementally and results in clusters of lipids with the same acyl chain composition (**Fig. 6b**). Within each cluster, the lipids are differentiated by their degree of unsaturation as the addition of a double bond decreases the CCS value almost linearly. Similar trends can be derived for triacylglyerides (**Fig. 6c**) and other lipid classes (**Suppl. Fig. 4**).

## DISCUSSION

Trapped ion mobility spectrometry is a particularly compact and efficient ion mobility setup, in which ions are held against an incoming gas flow and released as a function of their size and shape. The Bruker *timsTOF Pro* incorporates two TIMS analyzers in the front part of a high-resolution quadrupole TOF mass spectrometer and fully supports our recently introduced PASEF scan mode. In this study, we developed a nanoflow lipidomics workflow that takes full advantage of TIMS and PASEF.

In 100 ms TIMS scans, readily compatible with fast chromatography, we made use of the ion mobility separation of precursors to fragment on average nine per PASEF scan by rapidly switching the mass position of the quadrupole. Even though this translates into MS/MS acquisition rates close to 100 Hz, the ion count per spectrum, and thus the sensitivity, is determined by the TIMS accumulation time (here 100 ms or 10 Hz). In principle, this allows us to acquire MS/MS spectra for all detectable isotope patterns in short LC-MS runs, which would otherwise require much longer gradients or multiple injections and advanced acquisition strategies. While many of the acquired spectra remained unidentified in our current data analysis pipeline, the PASEF acquisition strategy generates very comprehensive digital MS/MS archives of all samples that can be mined with alternative and novel search algorithms at any time in the future.

Our results indicate that the nanoflow PASEF lipidomics workflow is readily applicable to a broad range of biological samples, such as body fluids, tissue samples and cell cultures. With a single extraction step, we quantified approximately 500 lipids with very high quantitative accuracy and reproducibility from as little as micrograms of tissue or only a few thousand cells. This makes our workflow very attractive for sample-limited applications, such as lipid analysis from biopsies, microdissected tissue or sorted cell populations.

An important application of lipidomics is the investigation of body fluids, for example blood plasma. Our analysis of a human reference sample was in good agreement with previous reports and surpassed the coverage for glycerol- and glycerophospholipids, using only a fraction of the analysis time and sample amount. Based on these results, we estimate a limit of detection in the attomole range for these analyte classes.

Our online PASEF lipidomics workflow positions each lipid in a four-dimensional space with a precision <2 ppm for masses, <0.2% for CCS and about 1-5% for retention times. We speculate that this precision and accuracy can be leveraged to facilitate lipid assignment in addition to the comprehensive MS/MS information generated by PASEF. To further explore this data space, we compiled a library of over 1,000 high-precision lipid CCS values directly from unfractionated biological samples. Our dataset largely extends the number of reported lipid CCS values and provides a basis for emerging machine learning techniques to predict CCS values more accurately and for a broader range of lipid classes.

We conclude that TIMS and PASEF enable highly sensitive and accurate lipidomics, and generate comprehensive digital archives of all detectable species along with very precise ion mobility measurements – a wealth of information which awaits full exploration and application. We also note that all the analytical advantages demonstrated for lipidomics should carry over to metabolomics in general, an area that we are currently exploring.

## METHODS

### Chemicals and biological samples

1-Butanol (BuOH), iso-propanol (IPA), ortho-phosphoric acid, formic acid, methanol (MeOH) and water were purchased from Fisher Scientific (Germany), and methyl tert-butyl ether (MTBE) from Sigma Aldrich (Germany) in analytical grade or higher purity. The human plasma reference standard NIST SRM 1950 was obtained from Sigma Aldrich. Human cancer cells (HeLa S3, ATCC) were cultured in Dulbecco’s modified Eagle’s medium (DMEM), with 10% fetal bovine serum, 20 mM glutamine and 1% penicillin-streptomycin (all from PAA Laboratories, Germany) and collected by centrifugation. The cell pellets were washed, frozen in liquid nitrogen and stored at −80 °C. Mouse liver was dissected from an individual male mouse (strain: C57BL/6) and snap frozen immediately. Animal experiments were performed according to the institutional regulations of the Max Planck Institute of Biochemistry and approved by the government agencies of Upper Bavaria.

### Lipid extraction

Lipids were extracted using an adapted MTBE protocol^46^. Plasma samples were thawed on ice and the sample preparation was performed at 4 °C. 200 µL cold MeOH were added to 1 μL of blood plasma and vortexed for 1 min. Subsequently, 800 μL of cold MTBE were added and the sample was mixed for another 6 min before adding 200 μL water. To separate the organic and aqueous phases, we centrifuged the mixture at 10,000 g for 10 min. The upper organic phase was collected and vacuum-centrifuged to dryness. To extract lipids from mouse liver, we first homogenized 1 mg of tissue in methanol and followed the extraction protocol described above. To extract lipids from approximately 5×10^5^ HeLa cells, they were lysed after addition of MTBE by sonication (Bioruptor, Diagenode, Belgium). The dried lipid extracts from all samples were reconstituted in BuOH:IPA:water in a ratio 8:23:69 (v/v/v) with 5 mM phosphoric acid for LC-MS analysis^17^.

### Liquid Chromatography

An Easy-nLC 1200 (Thermo Fisher Scientific) ultra-high pressure nano-flow chromatography system was used to separate lipids on an in-house reversed-phase column (20cm × 75μm i.d) with a pulled emitter tip, packed with 1.9 μm C_18_ material (Dr. Maisch, Ammerbuch-Entringen, Germany). The column compartment was heated to 60 °C and lipids were separated with a binary gradient at a constant flow rate of 400 nL/min. Mobile phases A and B were ACN:H_2_O 60:40% (v/v) and IPA:ACN 90:10% (v/v), both buffered with 0.1% formic acid and 10 mM ammonium formate. The 30 min LC-MS experiment started by ramping the mobile phase B from 1% to 30% within 3 min, then to 51% within 4 min and then every 5 min to 61%, 71% and 99%, where it was kept for 5 min and finally decreased to 1% within 1 min and held constant for 2 min to re-equilibrate the column. This gradient was extended proportionately for 60 min and 90 min experiments. We injected 1 μL in positive and 2 μL in negative ion mode on column and each sample was injected five times in both ionization modes.

### Trapped Ion Mobility – PASEF Mass Spectrometry

The nanoLC was coupled to a hybrid trapped ion mobility-quadrupole time-of-flight mass spectrometer (timsTOF Pro, Bruker Daltonics, Bremen, Germany) via a modified^19^ nano-electrospray ion source (Captive Spray, Bruker Daltonics). For a detailed description of the mass spectrometer, see refs. 36,37. Briefly, electrosprayed ions enter the first vacuum stage where they are deflected by 90° and accumulated in the front part of a dual TIMS analyzer. The TIMS tunnel consists of stacked electrodes printed on circuit boards with an inner diameter of 8 mm and a total length of 100 mm, to which an RF potential of 300 Vpp is applied to radially confine the ion cloud. After the initial accumulation step, ions are transferred to the second part of the TIMS analyzer for ion mobility analysis. In both parts, the RF voltage is superimposed by an electrical field gradient (EFG), such that ions in the tunnel are dragged by the incoming gas flow from the source and retained by the EFG at the same time. Ramping down the electrical field releases ions from the TIMS analyzer in order of their ion mobilities for QTOF mass analysis. The dual TIMS setup allows operating the system at 100% duty cycle, when accumulation and ramp times are kept equal^32^. Here, we set the accumulation and ramp time to 100 ms each and recorded mass spectra in the range from *m/z* 50 −1550 in both positive and negative electrospray ionization modes. Precursors for data-dependent acquisition were isolated within ±1 Th and fragmented with an ion mobility-dependent collision energy in the range from 25 eV to 63 eV. To improve the quality of the MS/MS spectra we acquired every PASEF scan twice, using a 50% higher collision energy in the second scan. The overall acquisition cycle of 1.2 s comprised two full TIMS-MS scan and ten PASEF MS/MS scans. Low-abundance precursor ions with an intensity above a threshold of 1,500 counts but below a target value of 10,000 counts were repeatedly scheduled and otherwise dynamically excluded for 0.4 min. TIMS ion charge control was set to 5e6. The TIMS dimension was calibrated linearly using four selected ions from the Agilent ESI LC/MS tuning mix [*m/z*, 1/K_0_: (322.0481, 0.7318 Vs cm^-2^), (622.0289, 0.9848 Vs cm^-2^), (922.0097, 1.1895 Vs cm^-2^), (1221,9906, 1.3820 Vs cm^-2^)] in positive mode and [*m/z*, 1/K_0_: (666.01879, 1.0371 Vs cm^-2^), (965.9996, 1.2255 Vs cm^-2^), (1265.9809, 1.3785 Vs cm^-2^)] in negative mode.

### Data analysis and bioinformatics

The mass spectrometry raw files were analyzed with MetaboScape version 4.0 (Bruker Daltonics). This version contains a novel feature finding algorithm (T-ReX 4D) that automatically extracts data from the four-dimensional space (*m/z*, retention time, ion mobility and intensity) and assigns MS/MS spectra to them. Masses were recalibrated with the lock masses *m/z* 622.028960 (positive mode) and *m/z* 666.019887 (negative mode). Feature detection was performed using an intensity threshold >800 counts in positive mode and 200 counts in the negative mode. The minimum peak length was set to 7 spectra, or 6 spectra in 3 out of 5 replicates when using ‘recursive feature extraction’.

Lipid annotation of the T-Rex 4D detected molecular features using MS/MS spectra was performed using the ‘high-throughput lipid search (HTP)’ function of SimLipid v. 6.05 software (PREMIER Biosoft, Palo Alto, USA). The lipid search comprised four lipid categories, Glycerolipids (GL), Glycerophospholipids (GP), Sphingolipids (SP) and Sterol lipids (SL) and TAG, DAG, PA, PC, PE, PG, PI, PS, Ceramides, Sphingomyelins, Neutral Glycosphingolipids, Steryl esters, Cholesterols and Derivatives, Oxidized glycerophospholipids classes. PE and PC lipids with ether- and plasmalogen-substituents were considered. Glycerophospholipids were only considered if containing an even number of carbons on at least one of the fatty acid chains. We searched for [M+H]^+^, [M+Na]^+^, and [M+NH_4_]^+^ ions in positive mode, and [M-H]^-^, [M+Cl]^-^, [M-CH_3_]^-^, [M+HCOO]^-^ and [M+AcO]^-^ in negative mode. The precursor ion and MS/MS fragment mass tolerances were both set to 10 ppm.

The initial search results were further filtered based on a ‘Matched Relative Intensity index’ >100 to ensure lipids are annotated based on MS/MS spectra with fragment ions corresponding to structure specific characteristic ions. The SimLipid results were manually inspected to remove potential false positives and to refine lipid annotations based on head-groups and/or fatty acyl composition as follows. We manually removed unlikely ion species e.g., [PC+NH_4_]^+^, and [PC+Na]^+^ if there were no detected fragment ions corresponding to neutral loss of the PC head group. Lipids from GP, ST, and SP categories were required to have their corresponding head group diagnostic ions observed e.g., *m/z* 369.3516 for cholesterol esters, *m/z* 184.073 for PC lipid species, and the neutral loss of 141 Th. TG/DG lipids with three/two unique fatty acid chains are reported only if at least two/one fatty acid chain fragment ion were/was detected. We report lipid identifications with (a) a short name (e.g. PC 32:1), (b) with a long name where the symbol @ indicates that this particular acyl-chain is not fully characterized by fragment ions (e.g. Cer d18:1_26:0@,), and (c) with a long name where head group and fatty acyl-chain composition are fully characterized by fragment ions (e.g. PG 16_1:16:1).

Lipid CCS values were predicted in MetaboScape based on a support vector machine learning approach by Zhou et al.^27^. Mass spectrometric metadata such as the PASEF frame MS/MS information were extracted from the .tdf files using the SQLite database viewer (SQLite Manager v3.10.1). Further data analysis and visualization was performed in Python (Jupyter Notebook) and Perseus (v1.6.0.8)^47^.

## Acknowledgements

We thank our colleagues in the department of Proteomics and Signal Transduction for discussion and help, in particular D. Voytik, A. Strasser, J. Mueller, I. Paron and C. Deiml. We acknowledge our colleagues H. Neuweger, N. Kessler and S. Wehner at Bruker Daltonics in Bremen for their support in data processing and analysis. This work was partially supported by the German Research Foundation (DFG– Gottfried Wilhelm Leibniz Prize), the Novo Nordisk Foundation (Grant agreement NNF14CC0001), the European Commission (FP7 GA ERC-2012-SyG_318987**–**ToPAG) and the Max-Planck Society for the Advancement of Science.

## Author contributions

C.G.V., K.S., M.M. and F.M. designed the research project; C.G.V., K.S., A.D.B. and F.M. performed the experiments; C.G.V., K.S., N.S.M., U.S.H., S.M., A.B. and F.M. analyzed the data; N.S.M. as well as U.S.H., S.M. and A.B. contributed analytical tools; C.G.V., M.M., F.M. wrote the paper.

