## Supplemental Information for "Trapped ion mobility spectrometry (TIMS) and parallel accumulation - serial fragmentation (PASEF) enable in-depth lipidomics from minimal sample amounts"

**Supplementary Tables**

**S1.** Lipids identified in human NIST SRM 1950 plasma extract. Lipids are grouped by the lipid class (column A). Further information such as retention time, *m/z*, ion mobility and CCS values, categorization, delta mass, number of double bonds, LSI ID, LIPID MAPS ID, matched fragment ions etc. are provided.

**S2.** Lipids identified in mouse liver extract. Lipids are grouped by the lipid class (column A). Further information such as retention time, *m/z*, ion mobility and CCS values, categorization, delta mass, number of double bonds, LSI ID, LIPID MAPS ID, matched fragment ions etc. are provided.

**S3.** Lipids identified in human HeLa cells extract. Lipids are grouped by the lipid class (column A). Further information such as retention time, *m/z*, ion mobility and CCS values, categorization, delta mass, number of double bonds, LSI ID, LIPID MAPS ID, matched fragment ions etc. are provided.

**S4.** Quantification (Intensity) values for lipids identified in human NIST SRM 1950 plasma extract. In the left table (orange), the number of 'zero' values has been calculated. In the right table, (green), the coefficient of variation (CV) has been calculated.

**S5**. Comparison of lipids identified in Quehenberger et al. study and our study. TRUE indicates the commonly identified lipids (color highlighted cells) based on the short name annotation while FALSE indicates lipids uniquely identified in one of the two studies.

**S6**. Comparison of lipids identified in Bowden et al. study and our study. TRUE indicates the commonly identified lipids (color highlighted cells) based on the short name annotation while FALSE indicates lipids uniquely identified in one of the two studies.

**S7.** CCS values across commonly identified lipids in the three biological samples (plasma, liver, HeLa) in the positive mode and their CV (column G).

**S8.** CCS values across commonly identified lipids (based on short name annotation) in our study compared with the Zhu and McLean laboratories.

**S9.** Experimentally and machine-learning predicted CCS values for 583 lipids identified in the positive mode of all three biological samples (plasma, liver, HeLa).

**S10.** Total 4D lipid dataset that comprises 1,327 CCS values of 926 unique lipids, representing the four major lipid categories and 15 lipid classes. Lipids are grouped by the lipid class (column A). Further information such as retention time, *m/z*, ion mobility and CCS values, categorization, delta mass, number of double bonds, LSI ID, LIPID MAPS ID, matched fragment ions etc. are provided. In the three last columns 'x' indicates the sample type extract where each particular lipid was identified.

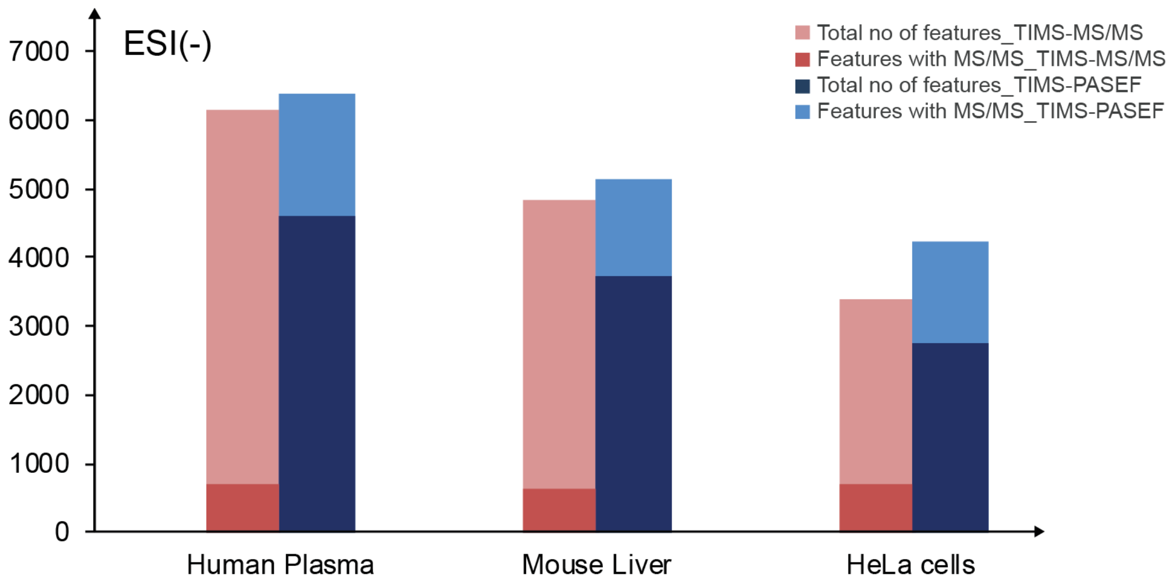


**Suppl. Fig.1**. **Evaluation of PASFE in lipidomics.** Total number of 4D features extracted from 30 min runs of human plasma (n=5), mouse liver (n=5) and human cancer cells (n=5) in negative ion mode without (TIMS-MS/MS, red) and with PASEF (PASEF, blue). The fraction of features assigned to MS/MS spectra is indicated by a darker color.

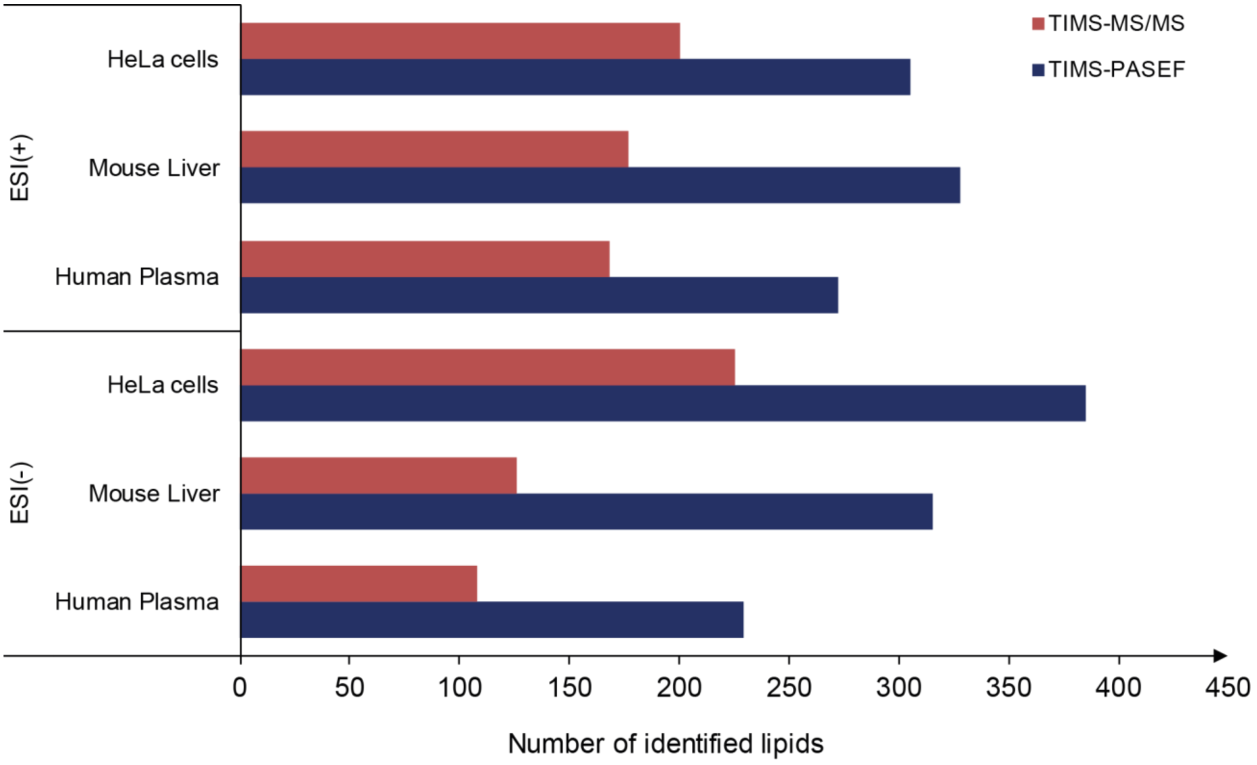


**Suppl. Fig.2. Lipid identifications in short LC-MS runs.** Total number of identified lipids in human plasma, mouse liver and human cancer cell extracts in both positive and negative ionization modes without (TIMS-MS/MS, red) and with PASEF (TIMS-PASEF, blue).

a.

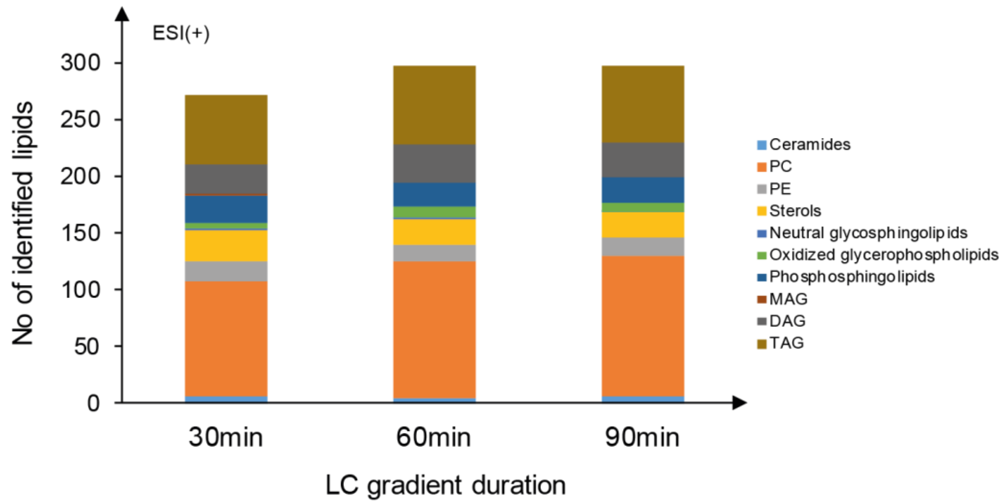


b.

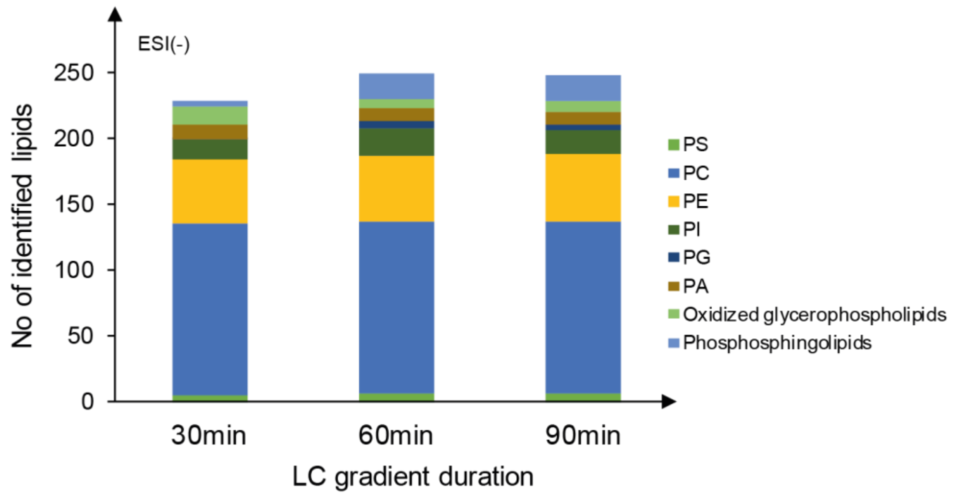


**Suppl. Fig.3. Lipid identifications with PASEF and increasing LC gradient duration. a,b,** Number of identified lipids from various lipid classes in 30, 60 and 90 min LC-MS runs of human plasma in **a,** positive and **b,** negative ionization mode.

*PC= Phospatidylcholine, PE=Phospatidylethanolamine, PA= Phosphatidic acid, PI= Phospatidylinositol, PG=Phospatidylglycerol, PS=Phospatidylserine, MAG=Monoacylglycerol, DAG=Diacylglycerol, TAG=Triacylglycerol.*

### Supplementary Figure 4

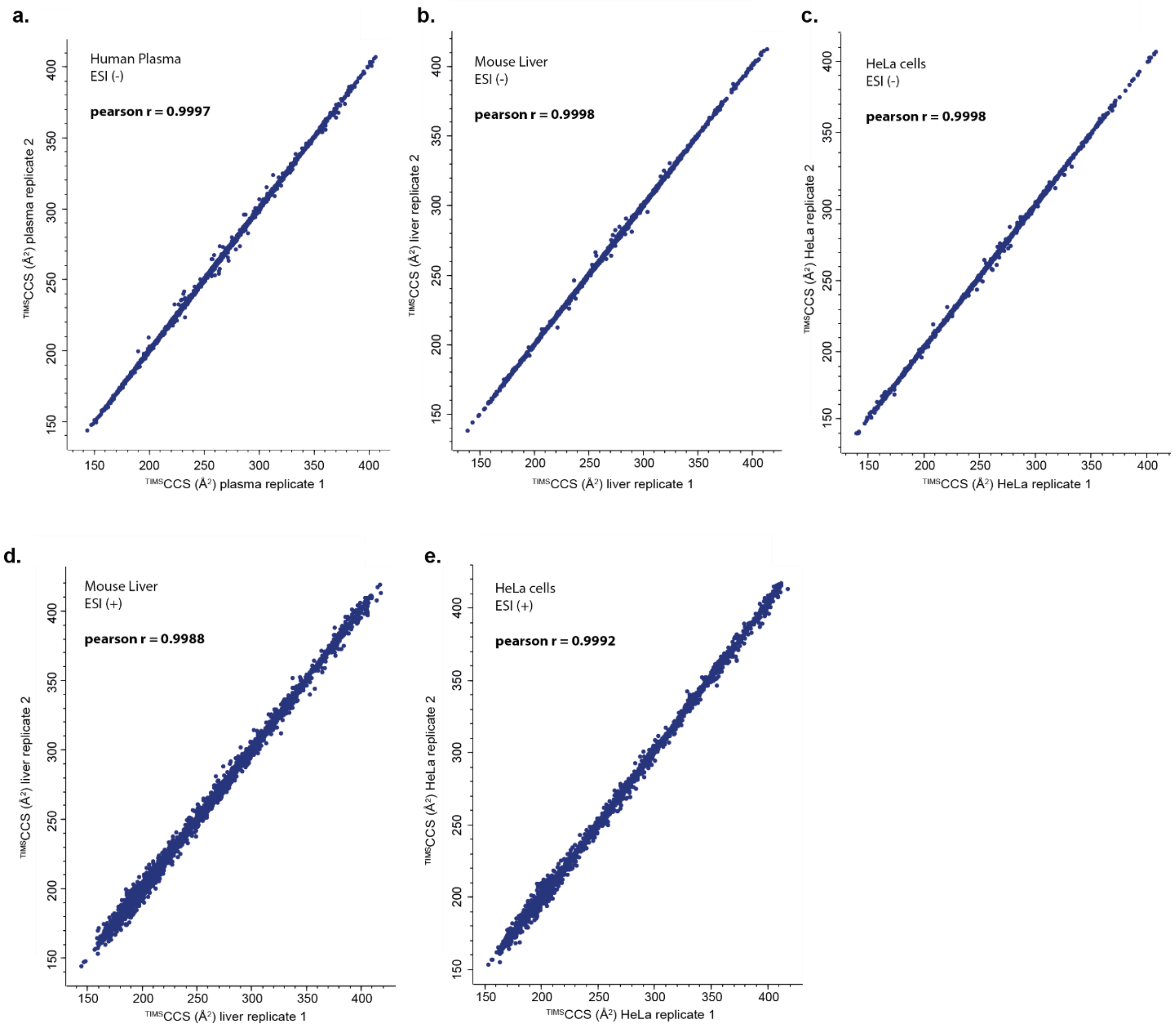


**Suppl. Fig.4. Precise measurement of $^{TIMS}$CCS values in lipid extracts from complex biological samples.** Pearson correlation of $^{TIMS}$CCS values of all 4D features detected in two replicate injections of **a,** human plasma extract in negative mode, **b,** mouse liver extract in negative mode, **c,** HeLa cells extract in negative mode, **d,** mouse liver extract in positive mode, **e,** HeLa cells extract in positive mode.

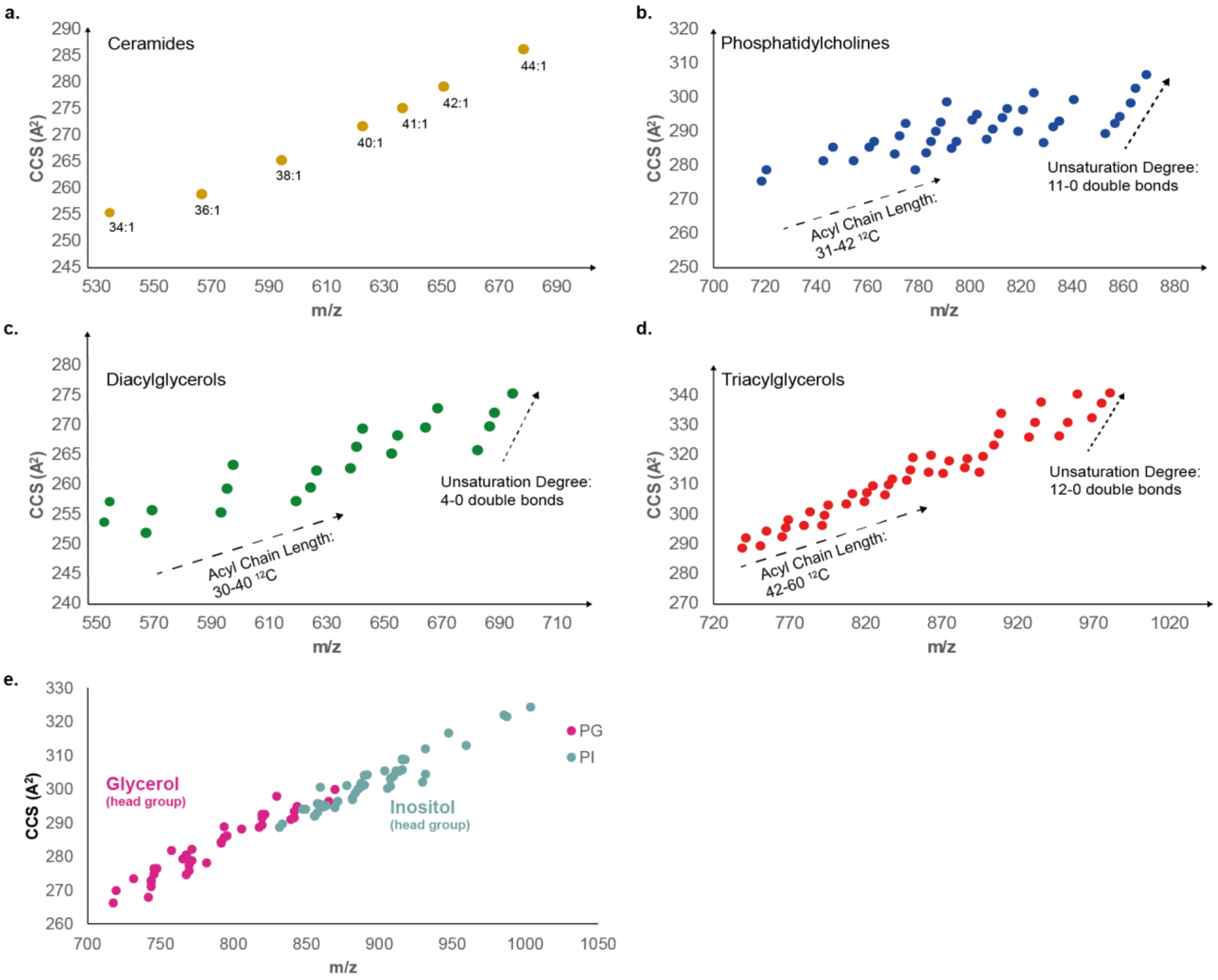


**Suppl. Fig. 5. Investigation of the effect of lipid composition on the CCS values. a-e,** Influence of acyl chain length and unsaturation degree on the CCS vs *m/z* distribution for **a,** Ceramides, **b,** Phosatidylcholines, **c,** Diacylglycerols, **d,** Triacylglycerols, and **e,** effect of head group (glycerol and inositol of glycerophospholipids).
